# MapToCleave: high-throughput profiling of microRNA biogenesis in living cells

**DOI:** 10.1101/2021.08.03.454879

**Authors:** Wenjing Kang, Bastian Fromm, Anna J. S. Houben, Eirik Høye, Daniela Bezdan, Carme Arnan, Kim Thrane, Michaela Asp, Rory B. Johnson, Inna Biryukova, Marc R. Friedländer

**Affiliations:** Science for Life Laboratory, Department of Molecular Biosciences, The Wenner-Gren Institute, Stockholm University, Stockholm, Sweden; UiT The Arctic University of Norway, Tromsø, Norway; Centre for Genomic Regulation (CRG), The Barcelona Institute for Science and Technology, Barcelona (BIST), Catalonia, Spain; Department of Tumor Biology, Oslo institute for Cancer Research, Oslo University Hospital, Norway; Analogs LLC, New York, USA; Department of Gene Technology, School of Engineering Sciences in Chemistry, Biotechnology and Health, KTH Royal Institute of Technology, Science for Life Laboratory, Solna, Sweden; Department of Medical Oncology, Inselspital, Bern University Hospital, University of Bern, Switzerland; Department for BioMedical Research, University of Bern, Bern, Switzerland; School of Biology and Environmental Science, University College Dublin, Ireland; Conway Institute for Biomolecular and Biomedical Research, University College Dublin, Ireland

**Keywords:** miRNA, microRNA, biogenesis, large-scale, comparative biology, RNA structure

## Abstract

Previous large-scale studies have uncovered many features that determine the processing of microRNA (miRNA) precursors, however, they have been conducted *in vitro*. Here we introduce MapToCleave, a new method to simultaneously profile processing of thousands of distinct RNA structures in living cells. Our new *in cell* method captures essentially all the biogenesis features that have been discovered through near two decades of in *vitro* studies - providing support for both approaches. We find that miRNA precursors with a stable lower basal stem are more efficiently processed and also have higher expression *in vivo* in tissues from twenty animal species. We systematically compare the importance of known and novel sequence and structural features and test biogenesis of miRNA precursors from ten animal and plant species in human cells. Lastly, we provide evidence that the GHG motif better predicts processing when defined as a structure rather than sequence motif, consistent with recent cryo-EM studies. In summary, we apply a new screening assay in living cells to reveal the importance of lower basal stem stability for miRNA processing and *in vivo* expression.

## Introduction

MicroRNAs (miRNAs) are short RNA molecules with important roles in animal gene regulation (Bartel, 2018). Since it has been estimated that mRNA from more than 60% of all human genes are regulated by miRNAs in one or more cellular contexts (Friedman et al., 2009), it is not surprising that these molecules have been found to play roles in biological processes, ranging from development (Giraldez et al., 2005) and formation of cell identity (Lim et al., 2005) to various diseases, including neurological illnesses and cancer (Esteller, 2011). Mutant animals that are completely devoid of miRNAs either die at early developmental stages (mice) or develop severe developmental defects (zebrafish) (Bernstein et al., 2003; Giraldez et al., 2005).

In the canonical biogenesis pathway, miRNA primary transcripts are transcribed by RNA polymerase II, often as molecules that are tens of thousands of nucleotides long (Cai et al., 2004; Lee et al., 2002, 2004). Each primary transcript harbors one or more hairpin fold-back structures, which are recognized by Drosha and its binding partner DGCR8 in the nucleus (Han et al., 2006). Drosha cleaves out the ~60 nucleotide long miRNA precursor, which is exported to the cytoplasm by Exportin-5 (Bohnsack et al., 2004; Lund et al., 2004; Okada et al., 2009; Yi et al., 2003). In the cytoplasm, the precursor is recognized and cleaved by Dicer, which is part of the canonical RNA interference pathway, thus releasing a ~22 nucleotide long RNA duplex (Bernstein et al., 2001; Hutvagner et al., 2001; Ketting et al., 2001; Knight and Bass, 2001). Subsequently, one of the strands of the duplex is selectively loaded into the Argonaute protein, which is a key component of the miRISC effector complex (Iwasaki et al., 2010). Once bound to Argonaute, the mature miRNA can guide the complex by partial base-complementarity to target mRNAs, which are then degraded through de-adenylation and de-capping or are translationally inhibited through obstruction of translation initiation (Bartel, 2009). There are numerous non-canonical miRNA biogenesis pathways (Ha and Kim, 2014); these all share the presence of a pre-miRNA hairpin structure and binding by an Argonaute effector protein promoting mRNA repression.

It has been estimated that the human genome harbors over 400,000 regions that could give rise to hairpins structures if transcribed (Bentwich et al., 2005). In contrast, the number of human precursors is estimated to be between 556 (Fromm et al., 2020) and 3000 (Friedlander et al., 2014), suggesting that the hairpins that enter miRNA biogenesis pathways are stringently selected. Many studies have evaluated hairpin features that license miRNA biogenesis. These assays have measured hairpin cleavage *in vitro*, testing numerous variants of a limited number of distinct hairpins (Auyeung et al., 2013; Fang and Bartel, 2015; Kwon et al., 2019; Li et al., 2020a). Through comparison of the variants that were processed and unprocessed, a number of structural features and sequence motifs have been identified. The overall structure with two single-stranded flanking sequences, a ~35 nucleotide double-stranded stem and a single-stranded apical loop is the key entry point into miRNA biogenesis (Fang and Bartel, 2015; Han et al., 2006). The sequence motifs UG at the basal junction, UGUG at the apical junction and CNNC at the 3’ flanking sequence have been reported to facilitate Drosha processing (Auyeung et al., 2013; Fang and Bartel, 2015). Recent studies have further found that miRNA precursors tend to have bulge-depleted regions in the upper and lower part of the miRNA duplex (Roden et al., 2017) and that bulges in the lower and middle part of the miRNA duplex influence Drosha processing efficiency and/or precision (Li et al., 2020b, 2020a). Other studies have shown that the GHG motif, defined as an unmatched nucleotide other than guanosine that is flanked by two base-paired guanosines at position −7 to −5 relative to the Drosha cleavage site, can facilitate miRNA precursor processing efficiency and precision (Fang and Bartel, 2015). However, there is some evidence that the GHG motif is better defined as a catalog of sequence/structure combinations (Kwon et al., 2019), and a recent cryo-EM study points to the importance of the structure itself (Jin et al., 2020).

Previous studies have been limited in that variants of only a few miRNA precursors have been tested, leaving open the possibility that some important biogenesis features may remain undiscovered. One recent study partly overcame this limitation by testing thousands of distinct RNA structures at the same time, providing evidence that structural uncertainty, measured as Shannon entropy, negatively influences processing (Rice et al., 2020). However, this experiment was conducted *in vitro*, so the contribution of the cellular context to miRNA biogenesis remains unstudied on a large scale.

Here we present MapToCleave, a novel method which can measure the processing of thousands of distinct RNA structures in living cells in a single experiment, recapitulating the details of natural miRNA biogenesis. Our approach is comparable to the one used by Chiang et al. (2010) (Chiang et al., 2010) to distinguish *bona fide* miRNAs from likely false annotations. We are expanding on this previous pioneering work to profile >10,000 structures in one experiment while Chiang et al. profiled up to ten structures per experiment. We find that miRNA precursors undergo differential processing in different cell types, underlining the importance of cell type-dependent processing. We also provide evidence that the precursors that are efficiently processed in our assay are significantly enriched in stable lower basal stem structures. We further extend this to *in vivo* conditions, showing that highly expressed miRNAs also tend to have stable lower basal stems in mammals, fruit flies and Lophotrochozoans – animal groups that are separated by >600 million years of evolution. Comparing the importance of known and novel features in predicting miRNA processing efficiency and *in vivo* expression, the lower basal stem ranks higher than several of the known sequence and structural motifs. Surprisingly, the known and novel features together explain only ~20% of miRNA processing. Lastly, we provide evidence that the GHG motif defined as a structure motif is a better predictor of miRNA processing efficiency and precision than is the motif defined as a sequence motif, supporting a recent cryo-EM study (Jin et al., 2020). In summary, our study extends the current model of miRNA biogenesis by revealing the lower basal stem to be an important structure that can tune miRNA processing and expression.

## Results

### MapToCleave measures processing of thousands of distinct RNA structures in cells

To systematically study miRNA biogenesis, we developed a novel high-throughput screening method - Massively Parallel Testing of Hairpin Cleavage (MapToCleave) - which we applied in a single experiment to simultaneously profile the processing of 12,472 distinct RNA structures in living cells. These structures include *bona fide* human miRNA precursors, non-human miRNA precursors and control non-hairpin sequences (Methods). First, the sequences were synthesized and cloned into an expression vector (Figure 1A, Methods). The generated expression constructs were pooled in a single library and then transfected into human cells, HEK-293T. The tested library was transiently expressed and the successfully transfected individual sequences were identified by DNA sequencing, while the structures that were successfully processed were detected by small RNA sequencing. By mapping the sequenced small RNAs back to the test structures, the biogenesis outcome of each RNA structure can be evaluated, as described below (Figure 1A, Supplemental Figure 1).

**Figure 1.**
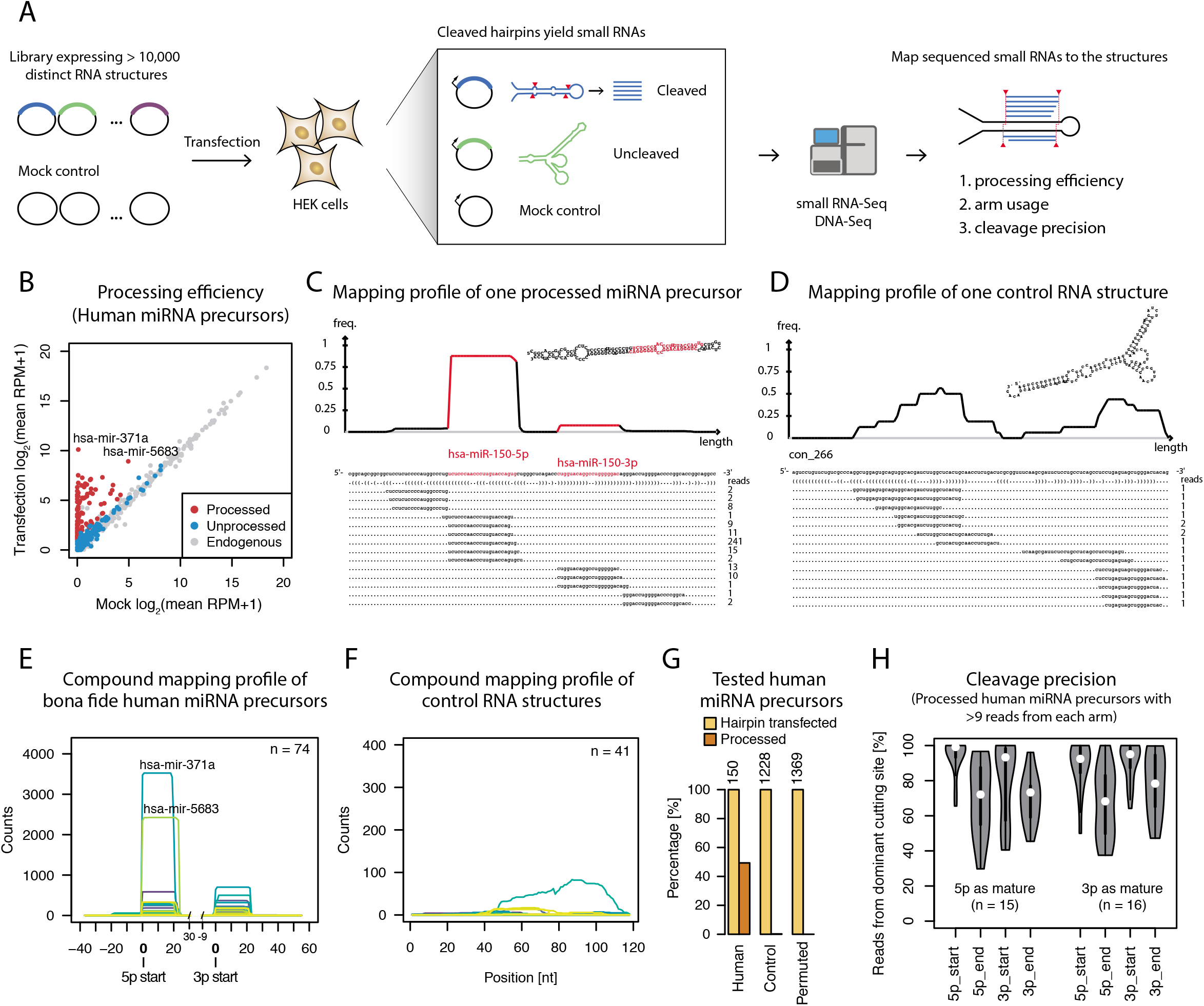
MapToCleave profiles miRNA processing of 12,472 distinct RNA structures. **(A)** Experimental design of MapToCleave. **(B)** Small RNA abundance in HEK-293T cells transfected with Mock controls or MapToCleave library (‘Transfection’). miRNAs that are part of the library and increase significantly in expression are defined as processed (red), while miRNAs that are part of the library but do not increase in expression are defined as unprocessed (blue). miRNAs that are endogenous to the cells and not included in the library are in grey. Expression is in log2 RPM (reads per million). **(C)** Example of MapToCleave processing of a *bona fide* human miRNA, showing clear patterns of Drosha and Dicer cleavage. A density plot of the read distribution of sequenced RNAs is shown above, and the exact read positions and read counts are shown below. **(D)** Example of a control non-hairpin RNA. The read profile is staggered, suggesting random degradation. **(E)** Compound read density plot of the 74 processed miRNA precursors. Each precursor is indicated with a distinct color. **(F)** Compound read density plot of 41 control non-hairpin RNAs, showing staggered patterns suggestive of random degradation. **(G)** Numbers of human miRNA precursors that are successfully transfected (yellow) and processed (orange). The same numbers are shown for control non-hairpin sequences from the human genome and control non-hairpins generated by randomizing (permuting) genome sequences. **(H)** MapToCleave processing precision of miRNA precursors. The assay recapitulates details of natural miRNA processing, including the increased precision of miRNA start positions relative to end positions.

Out of 150 *bona fide* human miRNA precursors successfully transfected into the HEK-293T cells (as measured by DNA sequencing), a total of 74 were efficiently induced and processed (Figure 1B, red dots, Methods). We found that the processing patterns of individual transfected precursors resembled known Drosha/Dicer processing signatures (Figure 1C), while the patterns for individual control sequences were staggered, suggesting random degradation (Figure 1D). This trend also holds when looking at compound distributions of read densities over the 74 processed human miRNA precursors (Figure 1E) and the 1228 control sequences (Figure 1F). Overall, while we found 49% of human miRNA precursors to be robustly cleaved in our assay, only 3/1228 (0.002%) control genomic non-hairpin sequences were processed, as were 0/1369 (0%) random non-hairpin sequences, showing the specificity of MapToCleave (Figure 1G). It is well established that miRNA strands tend to have more precise start than end positions (Czech et al., 2009; Khvorova et al., 2003; Okamura et al., 2009; Schwarz et al., 2003). We find the same tendency for miRNA strands in the MapToCleave library (Figure 1H), indicating that our high-throughput screening recapitulates subtleties of natural miRNA biogenesis.

### MapToCleave profiles cell-type dependent miRNA processing

A major advantage of MapToCleave is the ability to measure miRNA precursor processing in living cells, in the natural environment of protein cofactors, cellular compartments and more, in contrast to previous large-scale efforts to profile miRNA biogenesis which have all been *in vitro* (Auyeung et al., 2013; Fang and Bartel, 2015; Feng et al., 2011; Kwon et al., 2019; Li et al., 2020a; Nguyen et al., 2020; Rice et al., 2020). As a proof-of-principle, we tested human and murine MapToCleave precursors in human embryonic kidney HEK-293T cells and mouse NIH-3T3 fibroblast cells (Methods). In our replicate transfections in HEK-293T cells, we find only 3% (5/195) miRNA precursors to be differentially processed, showing the reproducibility of our method (Figure 2A, left). In contrast to these replicate experiments, when we compare processing in human HEK-293T versus mouse NIH-3T3 cells, we find that 16% (28/176) of the tested precursors are processed more efficiently in one of the two cell-types (Figure 2A, right). For instance, mir-872 is specific to the Glires animal group (rodents and lagomorphs) and is more efficiently processed in the mouse cell line (Figure 2A, right). Surprisingly, the human mir-130a also appears to be more efficiently processed in mouse cells than in human cells. Since this precursor appears to be more efficiently processed in other human cell lines (Supplemental Figure 2), this could be due to some specific blocking of this precursor in the HEK-293T cells. Based the difference in percentages between the replicate experiment (3%, above) and the between cell-type experiment (16%), we estimate that the biogenesis of 10-15% of mammalian miRNA precursors is substantially influenced by cell-specific factors. We consider this estimate to be a higher bound since we here change both the species and the cell type. In summary, we demonstrate that MapToCleave can profile cell type-dependent processing of miRNA precursors.

**Figure 2.**
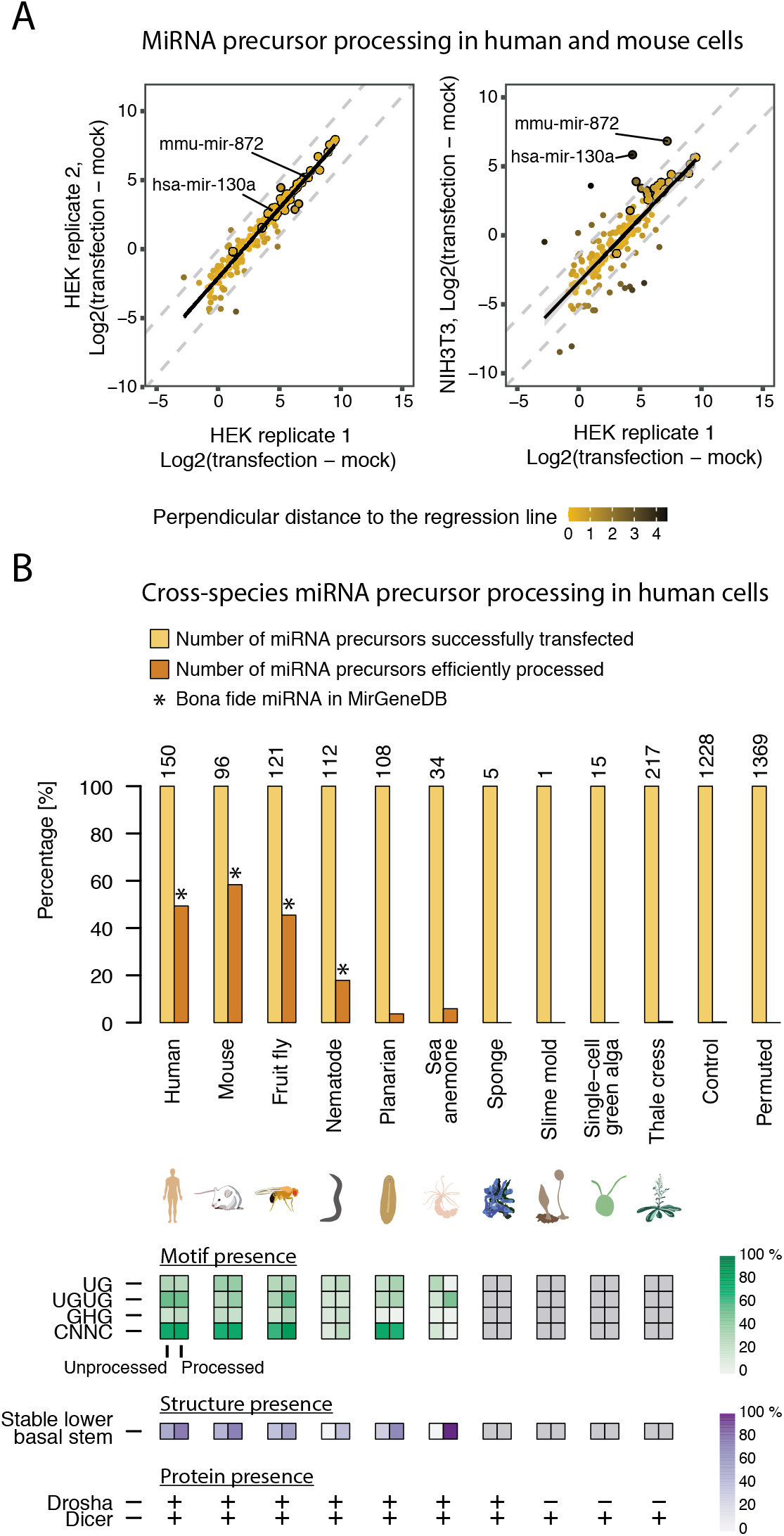
Cell-type dependent and cross-species miRNA precursor processing. **(A)** MapToCleave profiles cell-type dependent miRNA processing. The scatter plots show processing efficiency of the MapToCleave precursors in HEK-293T or NIH/3T3 cells with different transfection conditions. The processing efficiency is measured by the difference between mean precursor expression (RPM) in the transfection cells and in the mock cells. The color gradient from orange to black indicates the perpendicular distance from the dots to the fitted linear regression line. The high-confidence precursors with expression level higher than 5 RPM in the transfected cells are highlighted by a black circle. **(B)** Cross-species miRNA precursor processing in human cells. Above: Number of transfected (yellow) and number of processed (orange) precursors for 10 animal and plant species, and for control non-hairpin sequences. Below: Percentages of unprocessed and processed precursors that have sequence motifs known to facilitate processing (green). Also, percentages of unprocessed and processed precursors that have a stable lower basal stem structure (novel feature, in purple). Bottom: presence or absence of miRNA biogenesis factors in the 10 animal and plant species.

### miRNA biogenesis is functionally deeply conserved in animals

Having verified that MapToCleave recapitulates miRNA biogenesis in its natural cellular context, we next studied the processing of 709 non-human miRNA precursors in human HEK-293T cells to evaluate species-specific features of miRNA biogenesis. The precursors originate from species ranging from mouse, fruit fly, nematode, planarian, sea anemone, animal sponge, slime mold and single cell green algae to thale cress (*A. thaliana*) (Figure 2B). Given the evidence that miRNAs originated through convergent evolution in plants and animals (Axtell et al., 2011), it would be expected that phylogeny strongly affects biogenesis. Indeed, we find that the precursors from species that are more closely related to humans are more likely to be processed. The percentages of human, mouse and fruit fly precursors that are processed in human cells are comparable (ranging from 45-58%). In contrast, the percentage of nematode, planarian and sea anemone precursors that are processed in human cells is low (ranging from 4-18%). Sea anemone is the species most distant from humans in which we detect processing above trace levels, spanning a gap of >600 million years of evolution. This suggests that miRNA processing is deeply conserved, while the substrate preference varies in species as a function of phylogenetic distance. We find that essentially no animal sponge, slime mold, green algae or plant precursors are cleaved in human cells. miRNA biogenesis in animal sponges has previously been reported to be very different from other animals (Grimson et al., 2008), while slime mold and plant species do not have Drosha, which is a key biogenesis enzyme in animals (Avesson et al., 2012; Bologna and Voinnet, 2014; Bråte et al., 2018).

We next investigated if the observed processing efficiency can be explained by the known sequence motifs, which have been reported to facilitate mammalian miRNA processing. As has been previously reported, nematodes lack sequence motifs that are found in other animals (Auyeung et al., 2013), including planarians and sea anemone. However, there is no significant absence of sequence motifs in the precursors that are not processed in human cells (Figure 2B, green fields), suggesting that the low rate of processing has another explanation. Investigating the structures of the processed vs. unprocessed planarian and sea anemone precursors, we found the former had a tendency towards relatively structured and stable lower basal stems, defined as the first seven nucleotides of the double-stranded stem structure (Figure 2B, in purple, Supplemental Figure 3). This corresponds to positions −13 to − 7 relative to the Drosha processing site. This apparent importance of the lower basal stem suggests that it is worth revisiting the influence of structural features on miRNA biogenesis.

### Processed miRNA precursors have stable lower basal stems

We developed a new graphical representation to study the lower basal stem in more detail (Figure 3A). The ‘dumbbell’ heat plots show the structure of miRNA precursors, with the single-stranded region to the left and the apical loop to the right, the 5’ strand on top and the 3’ strand below. The color code indicates CG base pairing (dark blue), AU base pairing (light blue), GU base pairs (white) or bulges of mismatched nucleotides of increasing size (yellow to red). When summing precursors over human, mouse, fruit fly and nematodes, the most striking difference between the processed (Figure 3A, top) and the unprocessed miRNA precursors (Figure 3B, top) is at the lower basal stem from position −13 to −7 relative to the Drosha cleavage site (indicated with dotted white lines). Specifically, the precursors that are processed in our MapToCleave assay have fewer and smaller bulges in the lower basal stem (Figure 3A, top) than the precursors that are not processed (Figure 3B, top). This difference is observed in the ΔG minimum free energy estimates (Figure 3C, top) and is statistically significant (Figure 3D, top, p-value = 1.6e-12). We observe the same tendency when human (p-value = 0.014), mouse (p-value = 0.021), fruit fly (p-value = 0.020) and nematode (p-value = 0.001) precursors are studied separately, covering >600 million years of evolution. This tendency also holds true for the MapToCleave precursors tested in mouse cell lines (Supplemental Figure 4). To further support our findings, we re-analyzed miRNA precursor processing data from a previous study, in which Drosha cleavage efficiency of >50,000 sequence variants of three distinct primary miRNAs was tested *in vitro* in lysate containing Microprocessor (Fang and Bartel, 2015). By comparing the local structure profile of the variants with high, medium and low cleavage efficiency, we find that introducing a bulge at the basal stem has a more detrimental effect on Drosha processing compared to bulges in other regions (Supplemental Figure 5). In summary, we show that processed precursors have significantly more stable lower basal stem structures, from nematodes to human.

**Figure 3.**
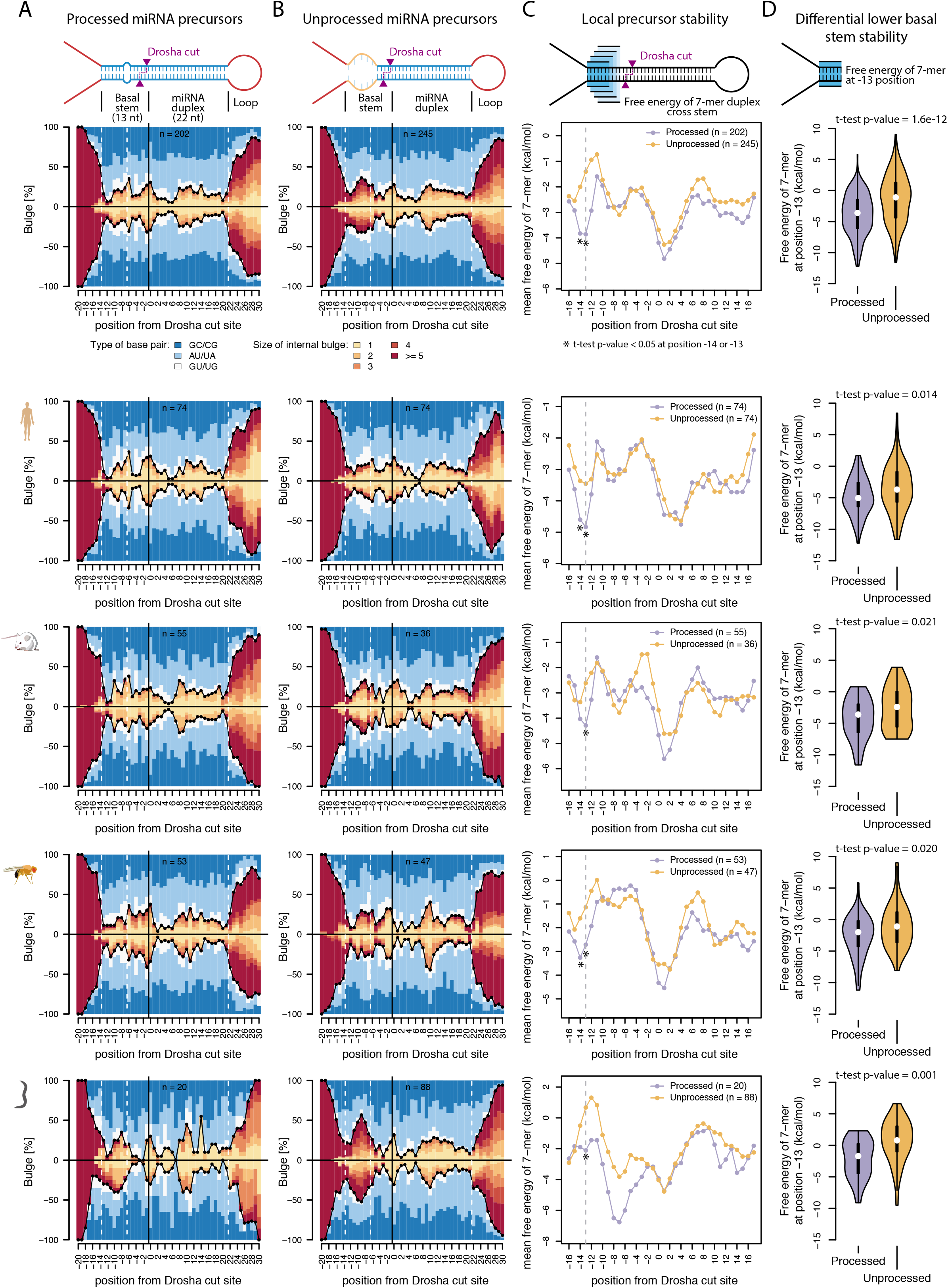
Processed miRNA precursors have more stable lower basal stem structures. Detailed structure profile of processed precursors **(A)** and unprocessed precursors **(B)**. The ‘dumbbell’ plots show the structure of miRNA precursors, with the single-stranded region to the left and the apical loop to the right, the 5’ strand on top and the 3’ strand below. The color code indicates CG base pairing (dark blue), AU base pairing (light blue), GU base pairs (white) or bulges of mismatched nucleotides of increasing size (yellow to red). The Drosha cleavage site at the 5’ strand is at position zero, and the two white vertical lines to the left indicate the position of the lower basal stem. **(C)** Thermodynamic stability profiles of processed and unprocessed precursors. The estimated minimum free energy (ΔG in kilocalories per mole) for RNA duplex was calculated by a rolling 7-nt window through the given precursor stem loop. Lower minimum free energy indicates more stable structures. **(D)** Minimum free energy distribution of the lower basal stem, represented by the 7-mer window at position −13, of processed and unprocessed precursors.

### Lower basal stem stability predicts miRNA expression levels *in vivo*

To test if the stable lower basal stem is a robust biological feature for miRNA processing rather than an artefact resulting from our MapToCleave screening system, we reanalyzed public small RNA sequence data from various animals. These data are from tissues and therefore represent *in vivo* expression, completely independent of our screening system. Specifically, we took advantage of the recently released second version of the manually curated microRNA gene database MirGeneDB (Fromm et al., 2020) and analyzed miRNA expression data composed of 191 tissue types from twenty species belonging to four clades: mammals, fruit flies, nematodes and lophotrochozoans. We averaged miRNA expression over tissues within a species and then compared the mostly highly and lowly expressed miRNAs within a given clade. By comparing the structure profile between the highly and lowly expressed miRNA precursors, we find that the lower basal stem is consistently observed to be more stable in the highly expressed miRNA precursors in mammals (p-value = 5e-5, Figure 4A-D, top row), fruit flies (p-value = 0.0084, second row) and lophotrochozoans (p-value = 0.013, fourth row). We do not observe the tendency in nematodes (p-value = 0.21, third row). It should be noted that nematode precursors have slightly longer basal stems (Warf et al., 2011), which in turn shifts the location of their lower basal stem (around from position −16 to −10) by around three nucleotides away from the Drosha cleavage site relative to the lower basal stem in other species (from position −13 to −7). Interestingly, the lower basal stem is also more stable in ancient miRNAs than in more recently emerged miRNAs (Supplemental Figure 6). These findings support the idea that the stable lower basal stem is not an artefact of our RNA structure screening system but is rather a naturally occurring and deeply conserved biological feature for miRNA processing.

**Figure 4.**
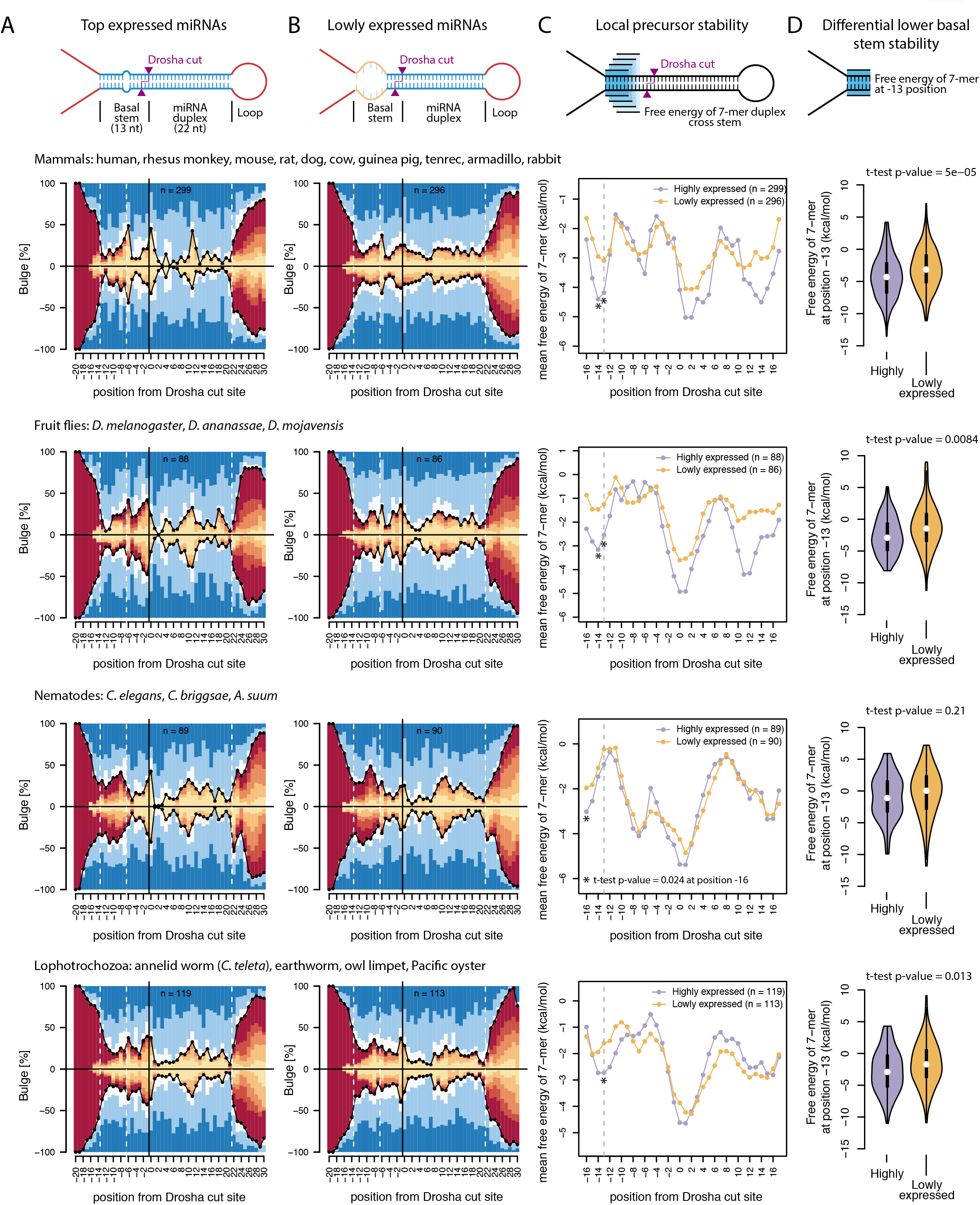
miRNAs with high *in vivo* expression have more stable lower basal stems. Similar to Figure 3, but the plots were generated based on the miRNAs with highest and lowest expression in animal tissues according to MirGeneDB.

### Chromatin-associated pri-miRNA profiles support importance of lower basal stem

Previous studies indicate that miRNA primary transcripts may stably associate with chromatin (Pawlicki and Steitz, 2008). To study whether precursors with stable and unstable lower basal stems give rise to different primary miRNA profiles as a result of processing, we reanalyzed sequenced primary miRNA transcripts associated with chromatin from a study by Conrad and colleagues (Conrad et al., 2014). In this previous experiment, the amount of intact vs. cleaved primary miRNA transcripts was used to estimate processing efficiency. We compared the structure profile between the most efficiently processed and the least efficiently processed miRNAs identified by the study (Supplementary Figure 7A-B). As expected, the efficiently processed miRNA precursors have a more stable lower basal stem compared to the non-efficiently processed miRNAs, though this tendency is only significant when counting from position −14 and not from position −13 (Supplementary Figure 7C-D). Again, this indicates the importance of the lower basal stem stability as a biological feature for miRNA processing.

### Design of miRNA precursors with improved or impaired processing capacity

Hairpin RNAs are widely used in RNA interference experiments and also for therapeutic treatments (Beg et al., 2017; Janssen et al., 2013; Sahu et al., 2019). We next investigated if it is possible to tune precursor design by modifying the lower basal stem regions. We designed four variants of mir-16, one of which should stabilize the lower basal stem and improve processing (variant 1) and three which should destabilize the lower basal stem and impair processing (variant 2-4, Figure 5A left and right panel). In each experiment we co-transfected with equimolar abundances of mir-30a and mir-125a for normalization, and all transfected miRNAs were additionally modified (tagged) in the mature region to discern them from endogenous miRNAs (Methods). We found that stabilizing the lower basal stem indeed improved expression subtly, while destabilizing the stem substantially reduced it (Figure 5A, middle, Figure 5B, Supplemental Figure 8). Interestingly, endogenous mir-99b, mir-501 and mir-1271 were consistently reduced in the transfection experiments (Figure 5B). These miRNAs may be part of the same regulatory networks as the three transfected miRNAs and may be repressed through negative feedback loops. The influence of stability of the lower basal stem can also be observed in the designed variants of mir-30a (Supplemental Figure 9). In summary, we show that hairpin design can be tuned by stabilizing or destabilizing the lower basal stem.

**Figure 5.**
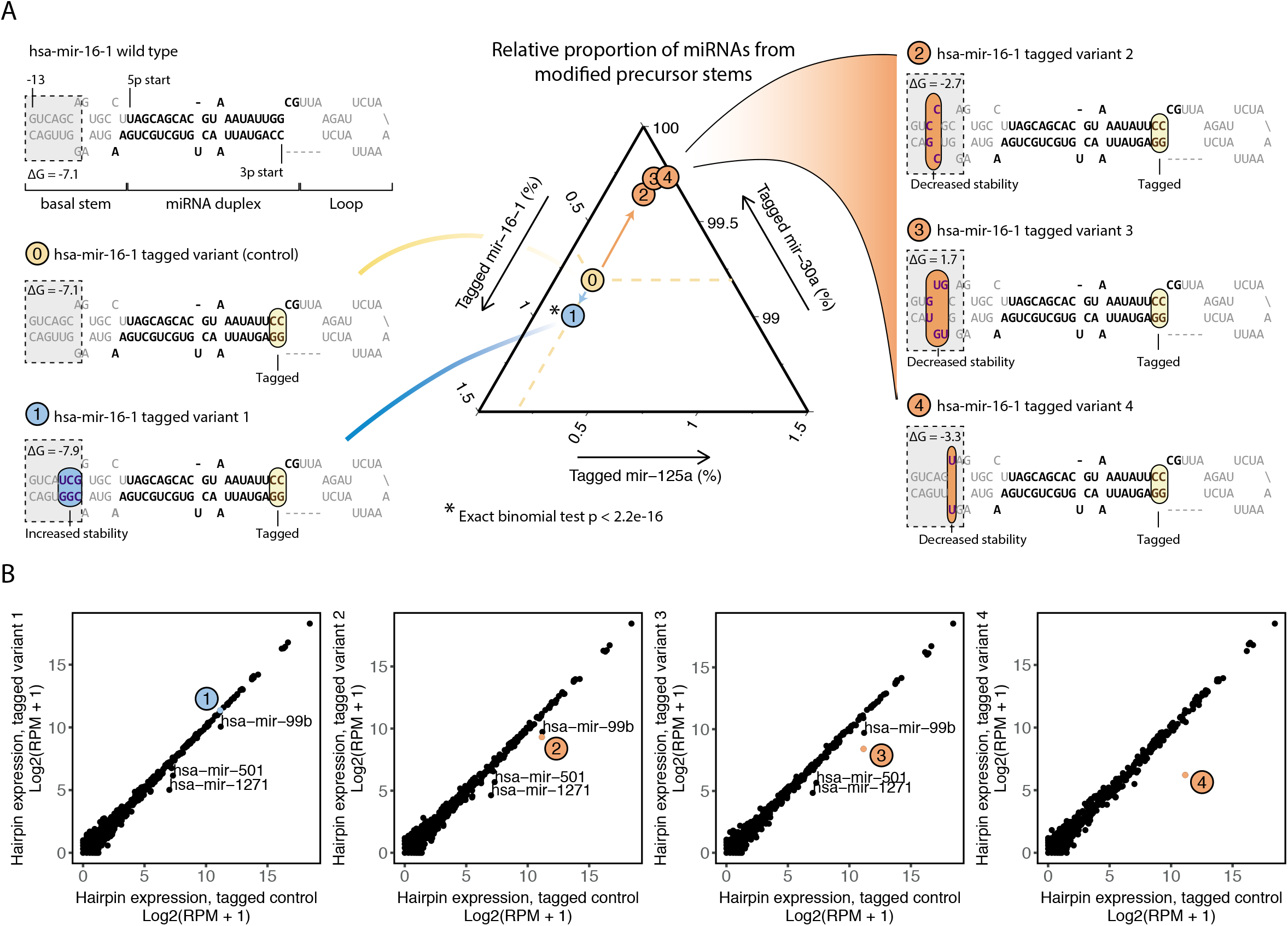
Design of miRNA precursors with improved or impaired processing. **(A)** Left and right panels: design of hsa-mir-16-1 variants with increased or decreased lower basal stem stability. All the variants are tagged by swapping two nucleotides at the 3’end of stem to distinguish them from endogenous miRNAs in sequencing. Middle panel: relative proportion of miRNAs from the tagged hairpin stem of hsa-mir-16-1, hsa-mir-30a and hsa-mir-125a, as measured by small RNA sequencing. **(B)** Scatter plots showing hairpin expression measured by summing 5p and 3p miRNAs in the mock or transfected cells.

### The GHG motif predicts processing better as a structure than as a sequence feature

Having focused on processing efficiency, we next investigated processing precision, measured as the percentage of sequenced miRNAs that map exactly to the consensus cut site (Figure 6A). We find that the precursors with high Drosha precision (>98%) tend to have a small bulge of one or two nucleotides that overlap with position −6, while the precursors that are processed with low precision (<90%) rarely have a bulge at this position (Figure 6B-C). This tendency for a bulge is clearly visible as an unstable region (Figure 6D). It is well-established that the GHG motif, located from nucleotides −7 to −5 from the Drosha cleavage site, can facilitate processing efficiency and precision of miRNA precursors (Fang and Bartel, 2015; Kwon et al., 2019). However, it is debated whether the motif is functionally more a sequence motif or a structural motif. Given the clear bulge that we see in precisely processed hairpins (Figure 6B), we propose the purely structural definition that a precursor has the GHG structure motif if it has a bulge composed of one or two nucleotides that overlap with position −6 (counted from 5’ stand). We find that the structural definition better predicts processing efficiency in our MapToCleave assay and also better predicts miRNA expression *in vivo* (Figure 6E, left) than does the sequence definition—that a precursor has a GHG motif if the positions −7 to −5 relative to the Drosha cleavage site consist of an unmatched nucleotide other than guanosine is flanked by two base paired guanosines (definition by Fang et al.) (Fang and Bartel, 2015). The same holds true for miRNA processing efficiency estimated from chromatin-associated miRNA primary transcripts from Conrad et al., 2014 (Figure 6E, middle). We also find that the structural GHG definition better predicts processing precision in the MapToCleave assay or *in vivo* (Figure 6E, right). In summary, we find that the GHG motif better predicts miRNA processing efficiency and precision when defined only by its structure.

**Figure 6.**
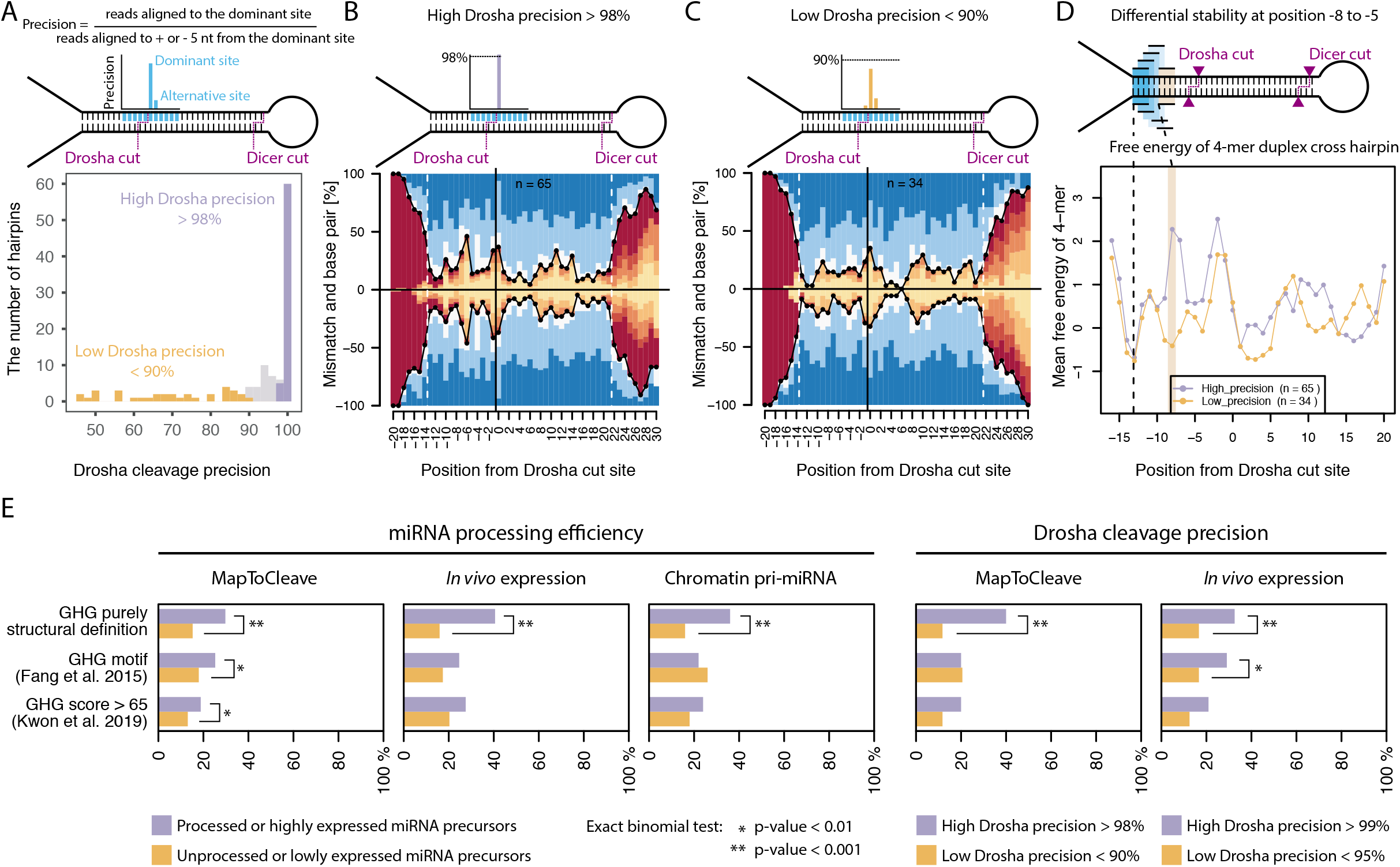
Influence of GHG feature on miRNA processing. **(A)** Histogram showing the Drosha cleavage precision of the processed precursors calculated by the equation on the top panel. **(B)** Detailed structure profile of precursors with high Drosha cleavage precision (>98% of reads from dominant cleavage site). **(C)** Same as **(B)** but using precursors with low Drosha cleavage precision (<90% of reads from dominant cleavage site). **(D)** Thermodynamic stability profile of the processed precursors with high and low Drosha cleavage precision. The free energy (ΔG in kilocalories per mole) was calculated by a rolling 4-nt window through the given precursor stem loop. The orange bar shows position −7. **(E)** The GHG motif predicts processing better when defined as a structural than sequence motif. miRNA precursors tested in our study were divided into the ones that are efficiently processed and highly expressed versus the ones that are unprocessed and had low expression. It was then tested how many miRNA precursors in the two groups that contained the GHG motif, according to three different definitions. The ‘GHG motif’ (Fang et al. 2015) and the ‘GHG score >65’ (Kwon et al. 2019) are defined by both structure and sequence features, while the ‘GHG structure’ is a purely structure feature. For the purpose of this analysis, the MapToCleave data from HEK-293T cells, MirGeneDB miRNA *in vivo* expression atlas of human tissues and chromatin associated pri-miRNA data from Conrad et al. 2014 were used.

### Relative importance of structures and sequence motifs for miRNA biogenesis

To understand the relative importance of known and novel sequence and structure features for miRNA biogenesis, we estimated how well each feature correlates with miRNA processing, as measured by MapToCleave, and miRNA *in vivo* expression, as collected in MirGeneDB (Figure 7A). Specifically, we applied linear regression to measure feature importance by the adjusted R squared value, which reflects the amount of data variance of miRNA processing efficiency that is explained by the model built on the feature (Methods). Intriguingly, the lower basal stem stability is ranked as the most important individual feature using MapToCleave data and the second most important using *in vivo* data (in green, Figure 7B), suggesting it is at least as important for processing as are the well-studied sequence motifs. We find that Shannon entropy (Rice et al., 2020) explains little of *in vivo* processing (in grey, Figure 7B), but does contribute to processing in our cleavage assay, although to a lesser extent than the lower basal stem stability (Figure 7B). Interestingly, two bulge-depleted regions of the precursors also contribute (in blue), consistent with previous results (Roden et al., 2017), as does the stability of other local structures along the miRNA stem that have only been investigated in a few studies (Li et al., 2020a; Nguyen et al., 2020). Overall, the combined structural features explain more of the miRNA processing (16.5%) than do the combined sequence features composed of CNNC, UG and UGUG (7.9%). The structural features explain comparable data variance of the *in vivo* expression (6.7%) to the sequence features (7.4%). In summary, we provide evidence that local structural precursor features are at least as important as the well-studied sequence motifs for miRNA processing.

**Figure 7.**
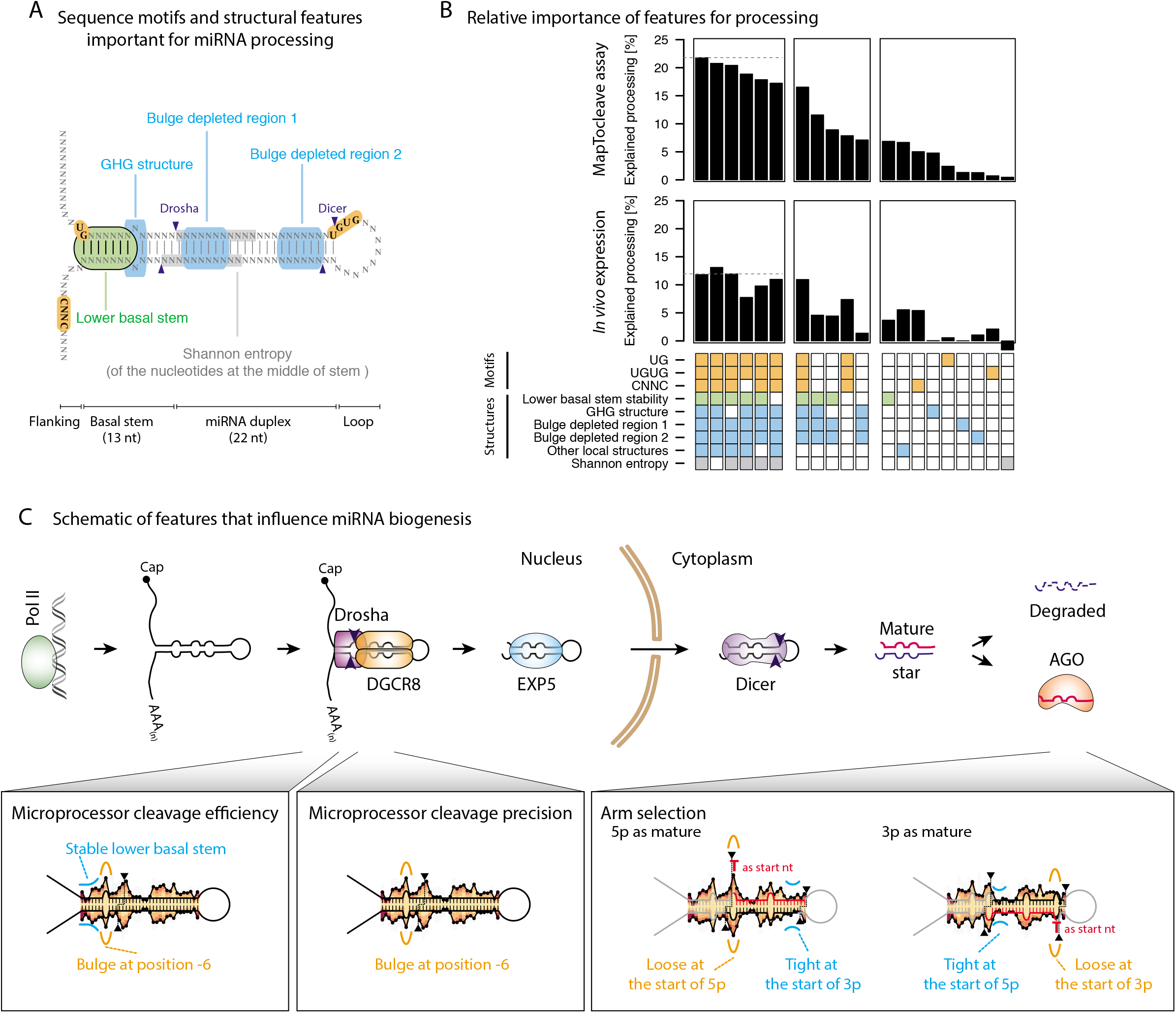
Relative importance of known and novel features for miRNA processing and expression. **(A)** Schematic of miRNA precursor stem showing location and type of known and newly identified features for miRNA processing efficiency. **(B)** Feature importance estimated by adjusted R squared value of the linear regression model with miRNA processing efficiency (MapToCleave data) or with mean miRNA RPM of human tissues (*in vivo* expression data from MirGeneDB) as outcome variable and a given feature (or features) as explanatory variable. **(C)** Schematic of features that influence miRNA biogenesis. The background structure profile in the panels of Microprocessor cleavage efficiency and precision shows the presence of bulges in the MapToCleave-processed precursors. The color code is the same as in Figure 3A. In the panel on arm selection, the background structure profiles on the left and right show, respectively, the presence of bulges in the 5p arm and the 3p arm selected MapToCleave processed precursors.

### MapToCleave recovers two rules of miRNA arm selection

Two rules have been proposed to determine which precursor arm gets selected as the guide miRNA and which gets degraded as a by-product of biogenesis (Czech et al., 2009; Khvorova et al., 2003; Okamura et al., 2009; Schwarz et al., 2003). According to the thermodynamics stability rule, the miRNA duplex end that is less stable is easier to open, and the arm whose 5’ end (so-called ‘5p’ arms) is at this end will be selected. According to the nucleotide rule, the arm with U and A as the first nucleotide is more likely to be selected as the guide miRNA compared to the arm with G and C. We divided the processed MapToCleave precursors into four groups depending on their preference for arm selection and investigated their distinct structural features (Supplemental Figure 10A). We find that precursors that have a strong 5p arm bias have a strong tendency for a bulge at the Drosha cleavage site (position 0), which would make the duplex end less stable, as predicted by the thermodynamic rule (Supplemental Figure 10B). Interestingly, for the precursors with a strong 3p bias, this bulge tends to be located at position −1, just outside of the duplex (Supplemental Figure 10A). The precursors that have a 3p arm bias also tend to have more bulges towards the 3’ end of the duplex (Supplemental Figure 10A), resulting in less stability in that end (Supplemental Figure 10B). Further, precursors with extreme 5p and 3p arm usage have the highest local free energy at the 5’ and 3’ end, respectively, of the miRNA duplex (Supplemental Figure 10B), and they also have the highest proportion of U and A, respectively, as the start nucleotide (Supplemental Figure 10C). These two rules of arm selection are identified by MapToCleave, suggesting the method is able to capture features that impact different steps of miRNA biogenesis (Figure 7C).

## Discussion

In this study, we have systematically surveyed features of miRNA biogenesis through the use of our high-throughput screening method MapToCleave. This allows us to test processing of thousands of distinct RNA structures in one experiment, recapitulating miRNA biogenesis in the natural context of living cells with protein cofactors, cellular compartments and more. We find that most of the tested human, mouse and fruit fly miRNA precursors are efficiently processed in human HEK-293T cells, while precursors of nematodes, planarians and non-bilaterian animals are inefficiently processed, and precursors of organisms that lack Drosha are not processed above trace levels (Figure 2B). Surprisingly, the miRNA precursors that are not processed in our MapToCleave assay specifically tend to have unstable lower basal stems, defined as positions −13 to −7 relative to the Drosha cleavage site (Figure 3). Applying public data of *in vivo* expression of curated miRNA complements of 20 animal species from MirGeneDB, we find that highly expressed miRNA precursors tend to have stable lower basal stems, while lowly expressed precursors tend to have unstable lower basal stems, indicating that the stability of this region tunes miRNA expression (Figure 4). We find that a structural definition of the GHG motif better predicts precursor cleavage efficiency and precision than does a sequence definition (Figure 6E), consistent with recent cryo-EM studies of Drosha substrate recognition (Jin et al., 2020; Partin et al., 2020). Comparing the relative importance of precursor features, we find that novel structural features explain MapToCleave processing efficiency and *in vivo* miRNA expression as well as or better than sequence motifs (Figure 7B). We find that lower basal stem stability in itself explains ~7% of processing efficiency—more than each of the individual known sequence motifs. Lastly, we recover and confirm known features of miRNA biogenesis, including the rules that determine miRNA strand selection (Supplemental Figure 10, Figure 7C).

Here we study miRNA biogenesis in cells and in tissues, which has the advantage that molecules are represented in natural concentrations and protein cofactors may be present. Methods applying purified Microprocessor have previously been widely used to study miRNA biogenesis *in vitro* (e.g. Han et al., 2006 (Han et al., 2006), Auyeung et al., 2013 (Auyeung et al., 2013), Fang et al., 2015 (Fang and Bartel, 2015), Li et al., 2020 (Li et al., 2020a), Rice et al., 2020 (Rice et al., 2020)). The *in vitro* approach has the advantage of specifically profiling Microprocessor activity without the confounding effects of nuclear transcription, export, Dicer processing etc. The two approaches seem complementary, and importantly the findings from our in-cell approach and in tissue approach converge and recover the main features that have previously been identified using *in vitro* systems (Figure 7B).

It may seem surprising that Shannon entropy explains little of in *vivo* miRNA processing (Figure 7B), in contrast to findings in a recent in *vitro* large-scale screening study (Rice et al., 2020). This may in part be explained by the complexity of living cells, but it may also be explained by the definition of miRNA precursors. The previous screening study used miRBase annotations, which contain many young miRNA genes as well as false positive annotations (Fromm et al., 2020). In contrast, our study uses MirGeneDB2 annotations, which are carefully curated. Thus, Shannon entropy may be a good measure for distinguishing genuine miRNAs from evolving genes or false positives (Supplemental Figure 11), while lower basal stem stability distinguishes genuine miRNAs that are highly or lowly expressed in tissues.

It is well-established that the length and stability of the ~35 nucleotide miRNA stem is important for processing (Fang and Bartel, 2015; Roden et al., 2017), and the contribution of the lower stem (positions −13 to −1 from the Drosha cleavage site) has been shown before in *in vitro* assays (Auyeung et al., 2013; Han et al., 2006; Zeng et al., 2005). Here we provide evidence that the first seven nucleotides of the lower stem (positions −13 to −7) are of particular importance relative to other individual sequence and structure features for miRNA expression in cells and in tissues (Figure 7B). We argue that this relates to Drosha recognition and binding, rather than simply defining the single-stranded to double-stranded transition, since the stability of the full seven nucleotides is critical and predicts processing much better than do shorter regions close to the single-stranded to double-stranded transition site (data not shown).

It may seem counterintuitive that the lower basal stem tunes miRNA expression, since a given precursor only gives rise to a single miRNA guide. However, there is evidence that many miRNA primary transcripts are not cleaved but rather remain relatively stable in the chromatin (Pawlicki and Steitz, 2008). Specifically, sequencing of RNAs in the chromatin allowed Conrad et al. to assign processing indexes to miRNA primary transcripts and to find that many had intermediate levels of processing (Conrad et al., 2014). If the lower basal stem facilitates efficient precursor processing, it would result in higher expression of the resulting mature miRNA, as we observe in the *in vivo* MirGeneDB data from 20 animal species.

Surprisingly, in our MapToCleave assay, we found that only ~50% of the *bona fide* human miRNA precursors were processed in HEK-293T cells. We estimate that ~5% of the tested precursors appear to be unprocessed because the exogenous expression is masked by high endogenous expression. We further estimate that ~9% of the tested precursors may not have been cleaved because they are normally clustered with other precursors that may facilitate their biogenesis (Fang and Bartel, 2020; Hutter et al., 2020; Kretov et al., 2020; Shang et al., 2020). The remaining unprocessed precursors tend to have unstable lower basal stems (Figure 3A), which means they may be outcompeted for Drosha processing by the precursors that have more stable lower basal stems or may have other structural features that facilitate interactions with Microprocessor. We did not find any depletion of the known sequence motifs in the unprocessed precursors (Figure 2B). Finally, it is possible that some of the precursors may depend on biogenesis cofactors that are absent in HEK-293T cells. This again highlights the advantage of studying miRNA biogenesis in a cellular system.

Interestingly, we find that known and novel precursor features overall explain less of the miRNA *in vivo* expression (13%) than they explain the miRNA processing (22%). This is what we expected, since MapToCleave comprises a well-controlled experiment in a human cell line, whereas the human MirGeneDB data comprises miRNA expression of various tissues that are affected by more layers of regulation of miRNA biogenesis as well as by the technical effects of heterogenous data. Even with our new features, the current model of miRNA biogenesis has a relatively limited information content and is still far from explaining the specificity of miRNA biogenesis. The optimal structure profile and the known sequence motifs together only explain ~22% of data variance of miRNA processing in MapToCleave (Figure 7B). Of the remaining ~78% data variance, MapToCleave DNA construct copy number for each precursor explains ~14%, consistent with previous findings that pri-miRNA transcription explains a substantial fraction of its final expression (de Rie et al., 2017). Besides data noise of experimental techniques, this points to more global factors, including, for example, RNA tertiary structure (Chaulk et al., 2011), global RNA structure (Rouleau et al., 2018), nuclear localization of precursors and biogenesis proteins, and biogenesis cofactors binding outside the local vicinity of the precursors (Nussbacher and Yeo, 2018; Treiber et al., 2017). Our results suggest that local features may only explain part of miRNA precursor selection and processing efficiency, and that a full model of miRNA biogenesis may also need to include global factors as critical components.

## Acknowledgements

We acknowledge the following funding sources: ERC Starting Grant 758397, “miRCell”; Swedish Research Council (VR) grant 2015-04611, “MapToCleave”; and funding from the Strategic Research Area (SFO) program of the Swedish Research Council through Stockholm University. RJ is supported by Science Foundation Ireland through Future Research Leaders award 18/FRL/6194. C. A. was supported by the Ministerio de Economía y Competitividad and FEDER funds under reference number BIO2011-26205 and BIO2015-70777-P and Secretaria d’Universitats i Investigació del Departament d’Economia i Coneixement de la Generalitat de Catalunya under award number 2014 SGR 1319. Anna J. S. Houben was funded as a Marie Curie post-doctoral fellow supported by the European Commission 7th Framework Program under grant agreement no. 330133. We thank Roderic Guigó, Xavier Estivill and Joakim Lundeberg for support and advice. The computations were enabled by resources in project [SNIC 2017/7-297] provided by the Swedish National Infrastructure for Computing (SNIC) at UPPMAX, partially funded by the Swedish Research Council through grant agreement no. 2018-05973.

## Author contributions

The study was conceptualized by M.R.F., I.B. and R.B.J. The MapToCleave library was designed by M.R.F. and R.B.J. and generated by A.J.S.H, D.B. and C.A. K.T. and M.A. prepared two sequencing libraries. All other experimental work was performed by I.B. Computational analyses were performed by W.K. under supervision of M.R.F., with analysis contributions from B.F. and E.H. The manuscript was written by W.K. and M.R.F., with contributions from all authors.

